# Intraspecific genetic and phenotypic diversity: parallel processes and correlated patterns?

**DOI:** 10.1101/288357

**Authors:** Fourtune Lisa, Prunier Jérôme G., Mathieu-Bégné Églantine, Canto Nicolas, Veyssière Charlotte, Loot Géraldine, Blanchet Simon

**Affiliations:** Centre National de la Recherche Scientifique (CNRS), Université Sabatier (UPS), UMR 5321 (Station d’Écologie Théorique et Expérimentale), Moulis, France; Université de Toulouse, UPS, UMR 5174 (Laboratoire Évolution & Diversité Biologique EDB), Toulouse, France; CNRS, UPS, École Nationale de Formation Agronomique (ENFA), UMR 5174 EDB, Toulouse, France

## Abstract

Intraspecific diversity plays a key role for evolutionary and ecological dynamics. It is the raw material on which acts selection, it improves species and communities resilience to disturbance and it affects the way species modulate their biotic and abiotic environment. Understanding patterns and underlying determinants of genetic and phenotypic intraspecific diversity is therefore of critical importance for ecological, evolutionary and conservation sciences. Here, focusing on two freshwater fish species (Gobio occitaniae and Phoxinus phoxinus) sampled across a large river basin (the Garonne-Dordogne river basin, France), we used causal analyses to test for genetic-phenotypic intraspecific diversity correlations (GPIDCs) and unravel the processes underlying intraspecific diversity patterns. Genetic diversity was assessed using microsatellite markers and phenotypic diversity was assessed through geometric morphometrics. We found disparities in the distribution of genetic and phenotypic diversity in the two species, suggesting higher level of local adaptation in G. occitaniae, and our results revealed common and contrasted processes shaping diversity at the α- and β-level. At the α-level, we found no GPIDC in both species despite common relations between isolation and genetic and phenotypic α-diversity in G. occitaniae. At the β-level, we found no GPIDC in P. phoxinus but we found a positive GPIDC in G. occitaniae. This correlation appeared to be caused by a direct impact of one facet of intraspecific diversity on the other, and we speculated that it could originate from positive assortative mating. Studying neutral genetic diversity and phenotypic diversity within an integrative framework appears as a valuable way of deciphering the complex and diverse impacts of neutral and adaptive processes on intraspecific diversity patterns.

## Introduction

In non-clonal species, all individuals are genetically and phenotypically unique, which constitutes the most elemental facet of biological diversity. Intraspecific biodiversity plays a key role in evolutionary and ecological dynamics (Bolnick et al. 2003, Odling-Smee et al. 2003). It is the raw material on which selection does act, potentially leading to adaptation to environmental changes, and improving population resilience to disturbances (Jung et al. 2013, Moran et al. 2015). Intraspecific diversity also affects the way populations modulate their biotic and abiotic environment, thus impacting community structure and ecosystem functioning (Hughes et al. 2008, Bolnick et al. 2011). Therefore, understanding patterns and underlying determinants of intraspecific diversity is of critical importance for ecological, evolutionary and conservation sciences (Chave 2013, Mimura et al. 2017).

As proposed for interspecific diversity, intraspecific diversity can be decomposed into two components: within-population (intraspecific α-diversity) and between-population intraspecific diversity (intraspecific β-diversity) (Loreau 2000). Within-population intraspecific diversity corresponds to the diversity space covered by individuals composing a population, whereas between-population intraspecific diversity corresponds to the differentiation observed among populations pairs. Intraspecific diversity also comprises a genetic and a phenotypic facet, the former being inherited from the parents and the later being affected by both inherited and non-inherited (environmental) information. Intraspecific genetic diversity is here defined as the variability of neutral and non-neutral genetic sequences observed within and among populations (Holderegger et al. 2006), whereas phenotypic diversity encompasses the diversity of individuals’ traits and includes behavioural, morphological and physiological traits (Violle et al. 2007).

Understanding how intraspecific diversity is maintained at the population level has long attracted ecologists and evolutionary biologists. For instance, the rise of molecular tools in the last decades has generated many studies describing patterns of intraspecific neutral genetic diversity (e.g. through allelic richness and FST), so as to unravel the demographic and evolutionary history of populations, and hence to improve their conservation and management (Manel et al. 2003, Reed and Frankham 2003, Blanchet et al. 2017). From an adaptive point of view, the relative importance of divergent natural selection in shaping the distribution of phenotypic traits across landscapes -and hence phenotypic β-diversity- has been the focus of studies combining quantitative genetics and experimental approaches (Kawecki and Ebert 2004, Leinonen et al. 2013, Blanquart et al. 2013). In parallel, ecologists have recently focused on the distribution of intraspecific phenotypic α-diversity across species and landscapes in order to better appraise its roles for community dynamics (Violle et al. 2012, Moran et al. 2015, Siefert et al. 2015). However, the study of intraspecific diversity still lacks an integrative framework in which patterns of genetic and phenotypic (α- and β-) diversity, as well as their underlying determinants, would be investigated simultaneously and considered as two potentially covarying facets of biological diversity. Remarkably, a framework in which two facets of biodiversity (namely species diversity and intraspecific genetic diversity) are studied in an integrative way has been introduced by Vellend (2005) and has generated an increasing number of studies (reviewed in Vellend et al. 2014, Lamy et al. 2017). These studies on species-genetic diversity correlations (SGDCs) led to a better understanding of the relationships between species and genetic diversity, as well as the processes shaping these facets of biodiversity in similar or contrasting ways (Taberlet et al. 2012, Vellend et al. 2014). Studying genetic-phenotypic intraspecific diversity correlations (GPIDCs) within a framework analogous to the SGDCs framework is attractive since genetic and phenotypic intraspecific diversity are intrinsically related and can be influenced by the same set of adaptive and neutral processes (Lowe et al. 2017). For instance, in the case of non-neutral genetic markers and adaptive traits, a positive GPIDC is expected when genetic and phenotypic non-neutral diversity are directly affected by environmental conditions (through selection and/or plasticity). In this case, both genetic and phenotypic α-diversity are expected to be high in populations inhabiting highly heterogeneous environments, and both genetic and phenotypic β-diversity are expected to be high between populations experiencing contrasting environmental conditions (Leimar 2005, Hedrick 2006, Wang and Bradburd 2014). Genetic and phenotypic diversity are also expected to be positively correlated if they are driven by neutral processes such as drift and dispersal (but see Edelaar et al. 2008, Lowe and McPeek 2014), which can notably be the case for neutral genetic markers and phenotypic traits that are weakly affected by selection (Hartl and Clark 2007). In that case, genetic and phenotypic α-diversity should be high in populations with large effective sizes and/or experiencing strong immigration. At the β-level, genetic and phenotypic diversity should be high between populations of small population sizes (Prunier et al. 2017) and/or geographically isolated from one another (Hutchison and Templeton 1999). Finally, a positive GPIDC can be explained by a direct relationship between genetic and phenotypic diversity, notably when genetic diversity directly codes for the considered traits, or appropriately describes the whole genomic diversity (Hoffman et al. 2014). Conversely, when genetic and phenotypic diversity are driven by (uncorrelated) divergent processes, GPIDCs are expected to be weak and non-significant.

Here, we aimed at testing spatial covariations in genetic and phenotypic intraspecific diversity in two parapatric species inhabiting a spatially-structured landscape, and at unravelling underlying determinants of each diversity facet at the landscape scale. More specifically, we first quantified and described genetic and phenotypic intraspecific diversity in two parapatric freshwater fish species (*Gobio occitaniae* and *Phoxinus phoxinus*) across an entire river drainage. We then investigated both α- and β-GPIDCs for these two species, and we finally deciphered the parallel or independent determinants shaping α- and β-genetic and phenotypic diversity using causal analyses (Fourtune et al. 2018). To this end, we gathered neutral genetic diversity and morphological diversity (a supposedly non-neutral type of trait) in both *G. occitaniae* and *P. phoxinus* so as to test whether or not the relative importance of main determinants of GPIDCs varied for species sharing a similar environment but with different life-history traits. We predicted that GPIDCs should be weak for the two species since neutral genetic diversity should mainly be driven by gene flow and/or drift, whereas morphology should be determined by local environmental characteristics. Alternatively, in river networks (Grant et al. 2007), factors affecting neutral processes (e.g. carrying capacity, geographic isolation, etc.) and adaptive processes (e.g. physico-chemical conditions such as water temperature, habitat heterogeneity, etc.) tend to covary along the network (e.g. upstream areas are generally homogeneous habitats with small carrying capacities whereas downstream areas are heterogeneous habitats with large carrying capacities) and these spatial covariation in underlying processes might generate strong GPIDCs. Moreover, the specific structure of river networks (treelike branching, constrained dispersal corridors, upstream-downstream environmental gradient) has already been theoretically and empirically shown to affect patterns of neutral and non-neutral diversity (Paz-Vinas and Blanchet 2015, Fronhofer and Altermatt 2017). Testing GPIDCs in highly spatially-structured landscapes such as dendritic riverine networks thus appears of particular interest.

## Materials and Methods

### Collection of genetic and phenotypic data

*Study species. Gobio occitaniae* (the Occitan gudgeon) and *Phoxinus phoxinus* (the European minnow) belong to the Cyprinidae family. Both species are insectivorous but differ in their foraging mode: *G. occitaniae* feeds predominantly on the bottom, whereas *P. phoxinus* feeds in the water column. *Gobio occitaniae* mean body length (120-150mm) is slightly larger than that of *P. phoxinus* (80-90mm). Moreover these parapatric species have contrasting levels of habitat specialisation: *G. occitaniae* lives in many habitat types and is ubiquitous in many river basins whereas *P. phoxinus* is more likely to occur in upstream (cold) areas.

### Study area and sampling

Fish were sampled across 48 sites evenly scattered across the Garonne-Dordogne river drainage (South-Western France). This river drainage covers a 79 800km^2^ area and sites were selected so as to cover the whole distribution of the two fish species, and hence their entire realized environmental niches. Electrofishing sampling was conducted during summers 2014 (42 sites) and 2015 (6 sites) and each site was visited once. Sampled area was of ∼500-1000 m2 to adequately represent the local habitat heterogeneity. *Gobio occitaniae* and *P. phoxinus* individuals were found in 39 and 34 sites respectively, with 25 sites in which the species were found in sympatry (see Fig. 1). We sampled up to thirty individuals per species and per sampling site (range, 21-30 and 24-30 for *G. occitaniae* and *P. phoxinus r*espectively, see Table S1), leading to the sampling of 1119 gudgeons and 978 minnows. Sampled individuals were anaesthetised using oil of clove before being carefully aligned on their right side on a white dashboard including a reference scale. The left side of each individual was photographed using a digital camera (Canon G16©) mounted on a tripod. Subsequently, we collected on each individual a small piece of pelvic fin which was preserved in 70% ethanol for genetic analyses. Individuals were then released alive in their respective sampling site.

**FIGURE 1.**
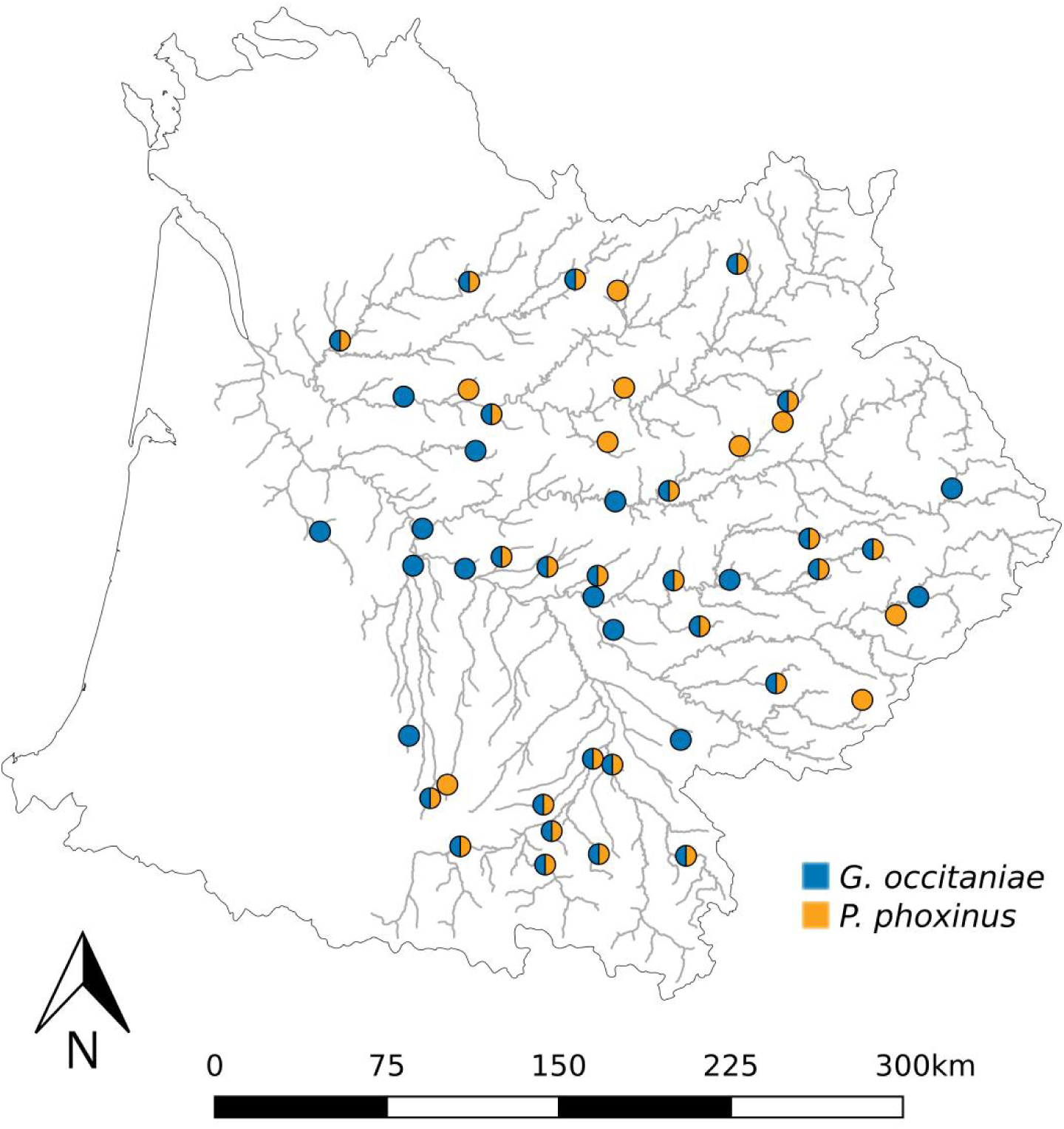
Location of the 48 sites sampled during summers 2014 and 2015 colored according to the species present.

**Table 1:**
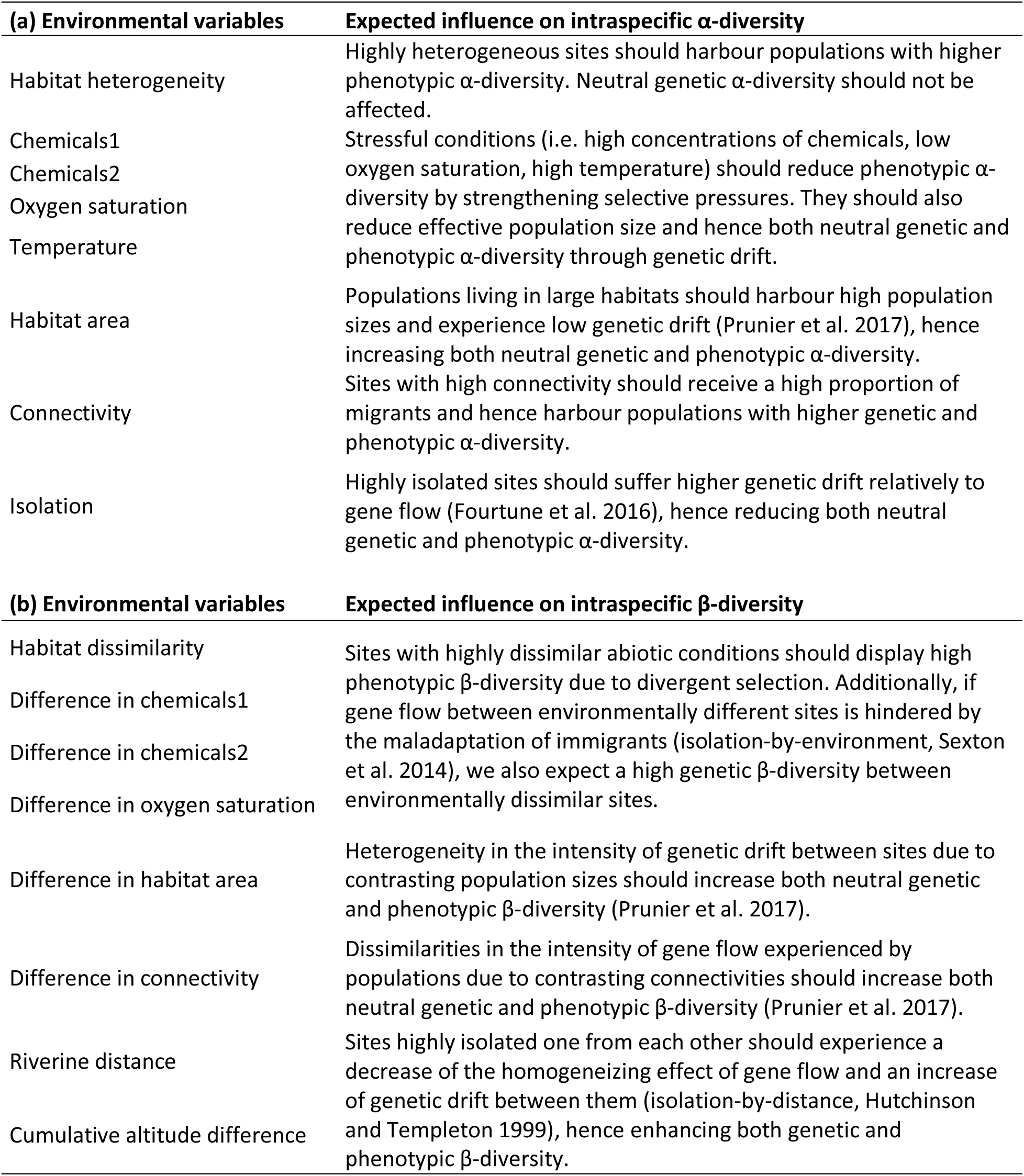
General predictions (and underlying processes) regarding the influence of environmental variables on intraspecidific α-diversity (a) and on intraspecidific β-diversity.

### Genetic data

Genetic DNA was extracted from all samples using a salt-extraction protocol (Aljanabi and Martinez 1997). Genotyping was performed using 15 and 18 microsatellite loci in *G. occitaniae* and *P. phoxinus* respectively. Accession numbers and conditions for polymerase chain reactions (PCR) are provided as supplementary material (Appendix S1). Genotypes were analysed using GENEMAPPER 5.0 (Applied Biosystems©). The presence of null alleles was assessed at each locus using MICROCHECKER 2.2.3 (Van Oosterhout et al. 2004). We also checked for gametic disequilibrium using GENEPOP 4.2.1 (Rousset 2008) after sequential Bonferroni correction to account for multiple tests. We discarded from further analyses any locus showing significant gametic disequilibrium and/or evidence of null alleles, leading to a total of 13 and 17 loci for *G. occitaniae* and *P. phoxinus* respectively (Appendix S1).

As a measure of genetic α-diversity, we computed -for each species-the allelic richness as the mean number of alleles across loci for a standardized sample size of 20 using ADZE 1.0 (Szpiech et al. 2008). As a measure of genetic β-diversity, we used three common indices based on allelic frequencies: Rousset’s linearized FST (FST/(1-FST)), hereafter denoted as FST (Rousset 1997), Nei’s version of Cavalli-Sforza’s chord distance *Da* (Nei et al. 1983) and Jost’s D (Jost 2008). Whatever the dataset, these three indices were highly correlated (Mantel r > 0.85, p <0.001): for the sake of simplicity, we thus only retained FST as a measure of genetic differentiation. This metric of genetic differentiation is indeed the most commonly used on population genetics and most theoretical works have been developed on this metric.

### Phenotypic data

Individuals morphology was analysed using a landmark-based geometric morphometrics approach (Rohlf and Marcus 1993). Sixteen homologous landmarks were defined so as to capture the overall body shape of each individual (Fig. 2). Landmarks coordinates were obtained from digitized pictures using the Pointpicker plugin (http://bigwww.epfl.ch/thevenaz/pointpicker/) in the ImageJ software (Schneider et al. 2012). As the distance between the camera and the fish slightly varied between sites, we size-corrected landmarks coordinates using the reference scale. For each species, landmarks were aligned using Generalized Procrustes Analysis (Rohlf and Slice 1990) with the R package *geomorph* (Adams and Otárola-Castillo 2013) in order to remove the effects of rotation, translation and scale on shape variation. Relative warps (n = 32) were computed for each individual by performing a Principal Component Analysis (PCA) on the aligned landmark coordinates of each species (Rohlf 1993). As the majority of the 32 relative warps explained a very small amount of variation, we only conserved for further analyses the first nine relative warps that together explained more than 85% of the variance in each data set (85.69% and 85.32% for *G. occitaniae* and *P. phoxinus* respectively). These relative warps were used as shape variables. Individual centroid size, which is the square root of the sum of squared distances from landmarks to their centroid, was used as a surrogate of overall body size of each individual (Bookstein 1991).

**FIGURE 2.**
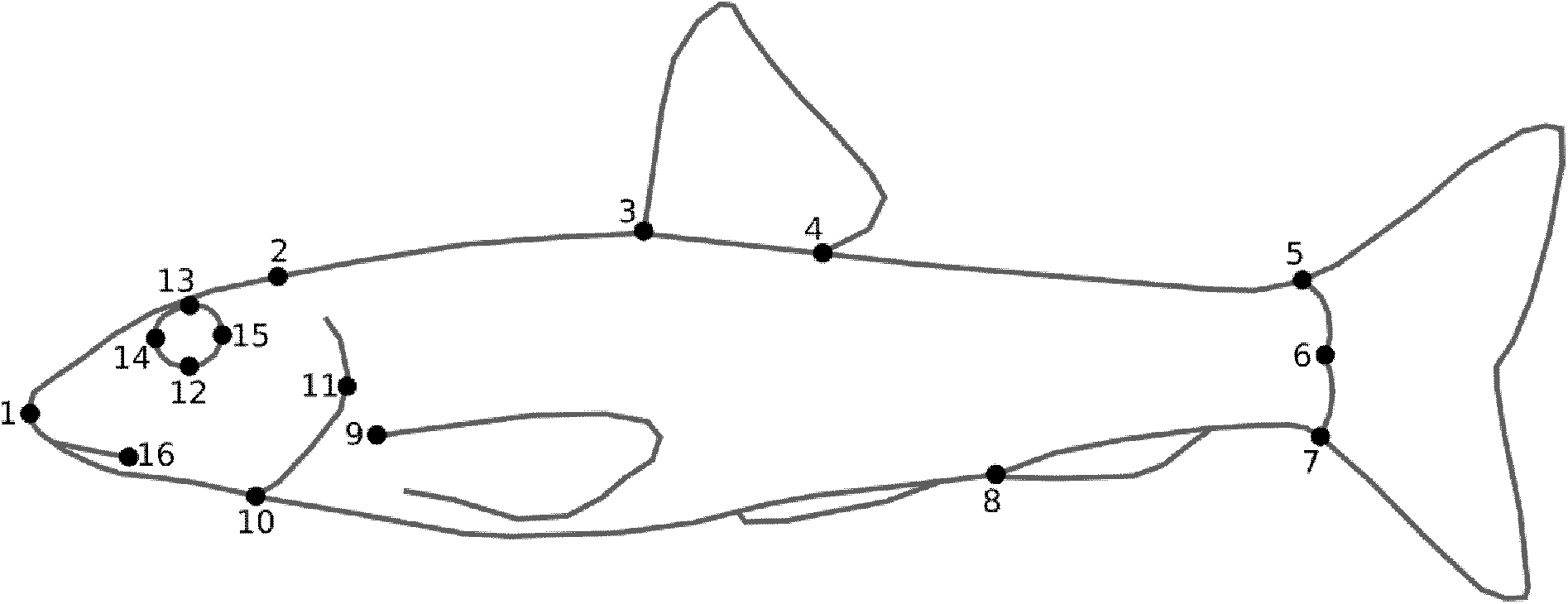
Location of 16 homologous landmarks used to assess phenotypic diversity in Gobio occitaniae and Phoxinus phoxinus. Landmarks refer to (1) tip of the snout, (2) beginning of scales coverage on the dorsal outline, (3) anterior and (4) posterior insertions of the dorsal fin, (5) dorsal insertion of the caudal fin, (6) posterior extremity of the body, (7) ventral insertion of the caudal fin, (8) anterior insertion of the anal fin, (9) superior insertion of the pectoral fin, (10) posterior border of the operculum, (11) posterior extremity of the operculum, (12) the inferior, (13) superior, (14) anterior and (15) posterior extremities of the orbital circumference, (16) posterior extremity of the premaxillar.

Since relative warps are PCA coordinates, they can be seen as coordinates in a nine dimensions shape space (i.e. the hyperspace in which each point represents a configuration of landmarks) of each individual. Morphological α-diversity was thus computed as the proportion of the total shape space (i.e. the shape space covered by all individuals from all populations for a given species) occupied by all individuals from a given population (see Fig. S2 for a graphical example), after accounting for differences in sampling sizes among populations using a random resampling approach. This index is equivalent to the functional richness index developed by Villéger et al. (2008) at the interspecific level. Morphological β-diversity was computed -for each species- as the euclidean distance between the consensus (i.e. the average shape in a population) of each populations pair (see Fig. S2). Additionally, in order to further decompose this index of differentiation, we computed the functional dissimilarity index Fβ initially developed by Villéger et al. (2011) for interspecific diversity and informing the proportion of the total shape space that is not shared between two populations from a given pair:

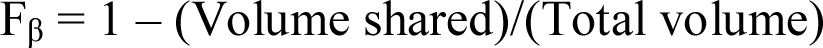

Fβ ranges from 0 to 1, with 0 indicating a perfect overlap in shape space occupation and 1 indicating that no space is shared between two populations. Intermediate Fβ can result from two (non-exclusive) mechanisms: turnover (the two populations fill distinct parts of the shape space with weak overlap) and nestedness (one population fills a small proportion of the shape space filled by the other) (Baselga 2010, Villéger et al. 2013). Consequently, we further computed Fpturn, the proportion of Fβ that is due to trait turnover so as to tease apart the effect of nestedness and turnover. Because of computing limitations, these two indices were computed only from the first three relative warps coordinates. All indices were computed using functions available at http://villeger.sebastien.free.fr/Rscripts.html.

### Collection of environmental data

We gathered several variables related to environmental characteristics and river topography. These variables are likely to impact intraspecific diversity through evolutive and/or neutral processes. General predictions related to the effect of each variable on intraspecific diversity are listed in Table 1.

### Environmental characteristics

*S*ubstrate type covering the river bed (i.e. the habitat for these fish species) was evaluated visually on each site following a predefined protocol: substrate was classified into nine categories based on particle size, ranging from silt (< 0.05mm) to solid bedrock (see Table S3), and the percentage of each category composing the river bed of each site was estimated visually within a predefined area of ∼100 m2 that was representative of the sampling area. From these data, habitat heterogeneity was computed as the Pielou’s evenness index, with low values identifying sites in which one of the substrate categories was dominant and hence relatively homogeneous habitats (large values indicated heterogeneous habitats with various substrate types). Habitat dissimilarities between sites were computed from percentages of substrate categories as Bray-Curtis distances, with a value of 1 identifying two sites sharing no substrate categories (i.e. sites highly dissimilar in substrate types). The other environmental variables were obtained for each site from the database of the Water Information System of the Adour Garonne basin (SIEAG, ‘Système d’Information sur l’Eau du Bassin Adour Garonne’; http://adour-garonne.eaufrance.fr) that gathers physico-chemical characteristics of surface water measured several times every year at numerous sites in the river catchment. Only sites for which data were available for July (a month in which the two species are highly active) of the years 2013, 2014 and 2015 were selected from the SIEAG database. The mean of the three values was calculated to inform the physico-chemical quality of the sites according to several parameters. We notably focused on two parameters directly affecting fish populations, i.e. oxygen saturation (%) (Crispo and Chapman 2008) and water temperature (°C) (Buisson et al. 2008). We gathered eleven additional variables informing overall water quality: concentrations in ammonium, azote, organic carbon, nitrate, nitrite, orthophosphate and phosphorus (mg/L), Biological Oxygen Demand (mg/L), water conductivity (mS/cm), pH and suspended matter (mg/L). We performed a PCA on these variables using R package “ade4” (Dray and Dufour 2007), and gathered the coordinates of each site on the first two axes (representing respectively 36.85% and 19.73% of the total variance) to create two synthetic variables (hereafter named *chemicals1* and *chemicals2*) informing water quality. High values of *chemicals1* correspond to high concentrations in ammonium, azote, organic carbon, phosphorus and a high Biological Oxygen Demand, whereas high values of *chemicals2* correspond to high concentrations in nitrate and nitrite and high values of conductivity, pH and suspended matter (see Fig. S4 for the graphical representation of the PCA).

**Table 3:** Coefficients estimates and significance obtained through full model averaging on the best (ΔAIC < 4) linear mixed-effects models (one per relative warp and per species) linking relative warps to the environmental variables and their associated quadratic terms models.(***: p-value < 0.001)

| Environmental variables | Relative warp 1<br>*Gobio occitaniae* | Relative warp 2<br>*Gobio occitaniae* | Relative warp 1<br>*Phoxinus phoxinus* | Relative warp 2<br>*Phoxinus phoxinus* |
| --- | --- | --- | --- | --- |
| Centroid size | 0.513*** |  | 0.455*** | -0.342*** |
| Centroid size<sup>2</sup> | -0.106*** |  |  |  |
| Connectivity |  |  | -0.017 |  |
| Distance from the mouth |  |  |  | 0.018 |
| Distance from the source |  |  | -0.025 |  |
| Habitat area |  |  | -0.016 |  |
| Slope |  | 0.009 |  | 0.134 |
| Carbone | 0.008 | -0.048 |  | -0.002 |
| Carbone<sup>2</sup> |  | -0.063 |  | 0.007 |
| Conductivity |  |  |  | -0.035 |
| Biological Oxygen Demand |  |  | 0.007 |  |
| Suspended matter |  |  |  | 0.013 |
| pH |  | 0.010 |  |  |
| pH<sup>2</sup> |  | 0.009 |  |  |
| Oxygen saturation | 0.272*** | 0.089 | 0.001 | 0.005 |
| Oxygen saturation<sup>2</sup> |  |  | 0.006 |  |
| Temperature | -0.236*** |  |  |  |
| Silt | -0.268*** | -0.016 |  |  |
| Cobble | -0.023 |  |  |  |
| Large cobble |  | 0.021 |  |  |
| Boulder |  | 0.007 |  |  |
| Large boulder |  | 0.045 |  | 0.080 |

### River topography

River distances from the outlet and from the source for each site, as well as river distance between each pair of sites, were computed using QuantumGIS software (QGIS; Quantum GIS Development Team 2017). Elevation for each site was obtained from the French Theoretical Hydrological Network (‘Réseau Hydrologique Théorique français’; Pella et al. 2012). A PCA was performed on elevation and distance from the outlet. The coordinates of each site on the first axis, accounting for 92.99% of the variance, were used to create a synthetic variable, hereafter named *isolation*, with high values corresponding to sites of high altitude, located far from the outlet and that are expected to be highly isolated geographically. Additionally, the cumulative altitude differences between each pair of sites along the riverine network were computed using the MATLAB software-coding environment (Mathworks, Inc., scripts available upon request). River width, used as a proxy for habitat area and hence carrying capacity (Raeymaekers et al. 2008), was characterised by measuring river bed width at two randomly selected locations for each sampling site, and subsequently computing the mean of these two values. The betweenness centrality value of each site was computed using ComplexNetGIS toolbox in ArcGIS (Caschili 2010). Betweenness centrality is an index quantifying the connectivity and positional importance of a node within a network (Freeman 1977, Estrada and Bodin 2008).

### Statistical analyses

Intraspecific genetic and phenotypic α- and β-diversities were compared between species using Wilcoxon rank sum test. Spearman rank correlations and Mantel tests were then used to assess and statistically test the significance of the correlations -for each species separately-between genetic and phenotypic diversity, at the α- and β-levels respectively.

The d-sep test (Shipley 2000, 2013) was used to unravel the relationships between environmental characteristics, topographical variables and intraspecific phenotypic and genetic diversity at the α- and β-levels. The d-sep test is a type of path (causal) analysis method computing the significance and likelihood of a causal model through the test of the conditional independences (named d-separations; Pearl and Verma 1987) that should be true if the model fits the data. A non-significant p-value associated with the null hypothesis that “the model fits the data” indicates that the observed data are consistent with the tested model. This method is very flexible as the statistical method used to test the independences is selected according to the data, which allowed in our case modelling both point summary (α-diversity) and pairwise (β-diversity) data types (Fourtune et al. 2018). Prior to analyses, environmental variables were log-transformed if needed to obtain a normal distribution, and all variables were centred to the mean and scaled.

At the α-level, we defined a causal model in which intraspecific genetic and phenotypic diversity were both linked one to the other and linked to oxygen saturation, water temperature, habitat heterogeneity, *chemicals1*, *chemicals2*, connectivity, *isolation*, and habitat area (see Table 1 for specific predictions). As some of the topological and environmental variables are expected to covary spatially, paths taking into account these covariations were included when needed. This model was tested using a d-sep test in which d-separations (i.e. path coefficients) were tested using linear regressions. This model was then simplified by removing paths one by one until reaching the model with the lowest Akaike Information Criteria (hereafter AIC; Burnham and Anderson 2002) so as to identify the main determinants underlying phenotypic and genetic α-diversity.

For genetic and phenotypic β-diversity, four environmental variables (oxygen saturation, temperature, *chemicals1* and *chemicals2*) were converted into pairwise environmental differences between sites using euclidean distances. Differences in habitat area, used as a proxy for carrying capacity and hence for the effect of genetic drift, were computed as *di* (distance based on the inverse; Relethford 1991) as recommended in Prunier et al. (2017). Additionally, we considered the three variables already taking the form of pairwise matrices: topographic distances, differences in cumulative altitude and habitat dissimilarities between sites. We defined a model in which genetic and phenotypic β-diversity were both linked one to the other and linked to these eight explanatory variables. A full model was tested using a d-sep test procedure recently developed for handling pairwise matrices (Fourtune et al. 2018) and that uses permutations-based linear regressions. This model was simplified using the same procedure as above, until reaching the model with the lowest AIC score.

As a side objective aiming at better understanding the spatial distribution of phenotypic diversity in the two fish species, we investigated phenotype-environment relationships by assessing and testing relationships between the individual shape of fish and raw environmental variables. For the sake of clarity, only the first two relative warps (respectively encompassing 31.3% and 15.5% of the variance in *G. occitaniae* and 27% and 21.9% of the variance in *P. phoxinus*) were separately considered in this analysis combining model selection and model averaging. Global models (one per relative warp and per species) linking relative warps to the environmental variables and their associated quadratic terms were implemented using the *lme* function in R package ‘nlme’ (Pinheiro et al. 2016) with the population identity included as a random-intercept effect. We also added individual centroid size and its quadratic term as explanatory variables to take the effects of allometry into account (Outomuro and Johansson 2017). All possible models were generated from the global model and their AIC were computed using the *dredge* function in the R package ‘MuMIn’ (Bartoń 2016). Full model averaging was then applied across the best models (ΔAIC < 4; Burnham and Anderson 2002) with the function *model.avg* in order to estimate the relative importance of each explanatory variable and weighted estimates associated to explanatory variables. All statistical analyses were performed with the R software (R Development Core Team 2017).

## Results

### Alpha- and β-intraspecific diversity

Both genetic and phenotypic α-diversity were higher for *P. phoxinus* than for *G. occitaniae* (Wilcoxon rank sum tests, W = 149, P < 0.001 for genetic α-diversity; W = 340, P < 0.001 for phenotypic α-diversity; Fig. 3a and 3b), indicating that minnow populations were on average more diverse genetically and phenotypically than gudgeon populations. However, within-species, we did not find significant correlations between genetic and phenotypic α- diversity (i.e. α-GPIDCs) for any of the two species (Spearman rank correlation tests, ρ = 0.105, P = 0.521 in *G. occitaniae* and ρ = 0.016, P = 0.927 in *P. phoxinus*, Fig. 4a and 4b).

**FIGURE 3.**
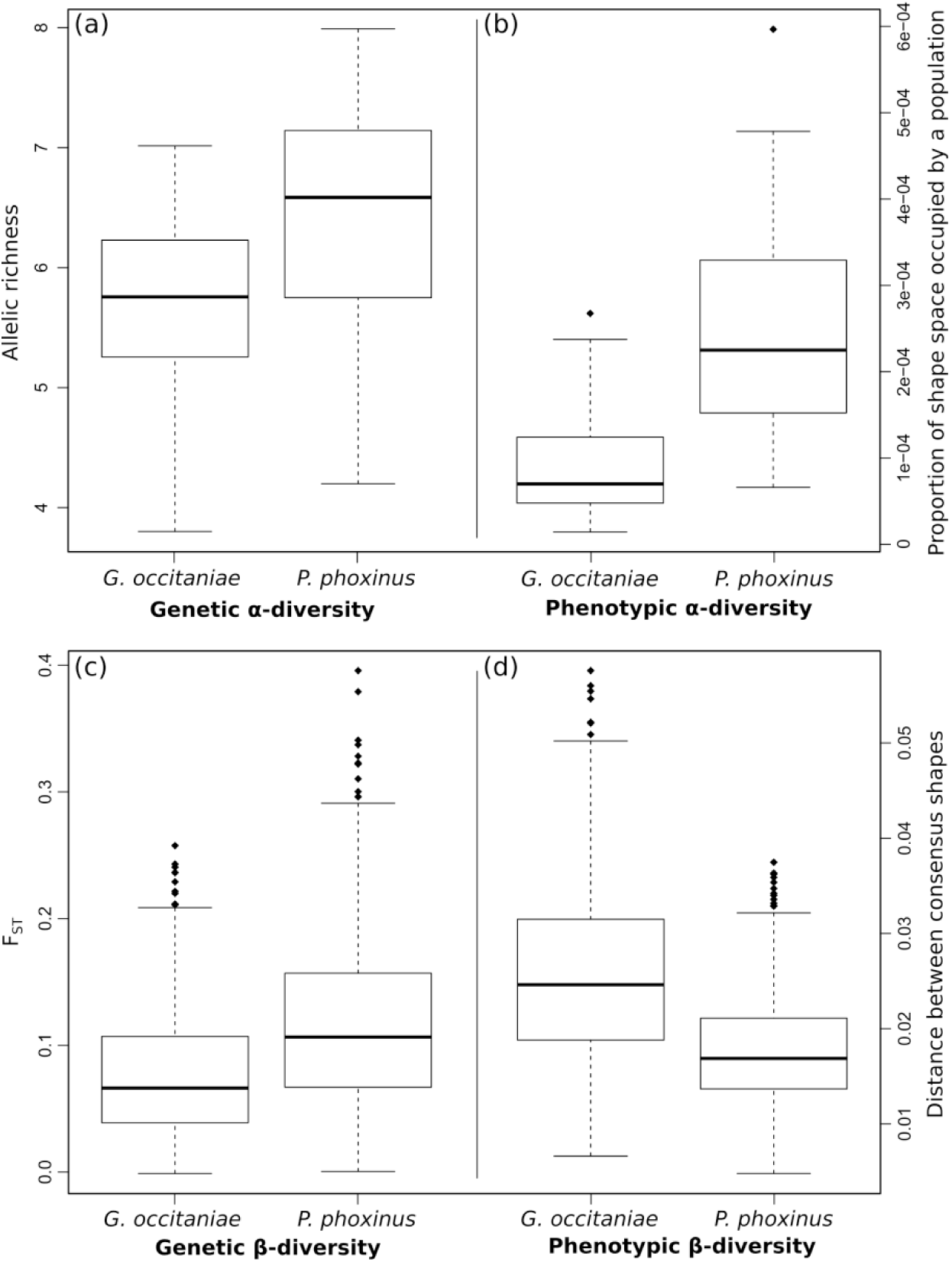
Boxplots summarizing the genetic α-diversity (allelic richness) (a), phenotypic α-diversity (proportion of shape space occupied by each population) (b), genetic β-diversity (FST) (c) and phenotypic β-diversity (euclidean distance between the consensus shapes of each pair of populations) (d) in *Gobio occitaniae* and *Phoxinus phoxinus*. The solid line within each box marks the median; the length of the box is the interquartile range (from the first to the third quartile). The lower whisker extends to the first quartile minus 1.5 times the interquartile range; the upper whisker extends to the third quartile plus 1.5 times the interquartile range. Diamonds represent the data points which are beyond the whiskers.

**FIGURE 4.**
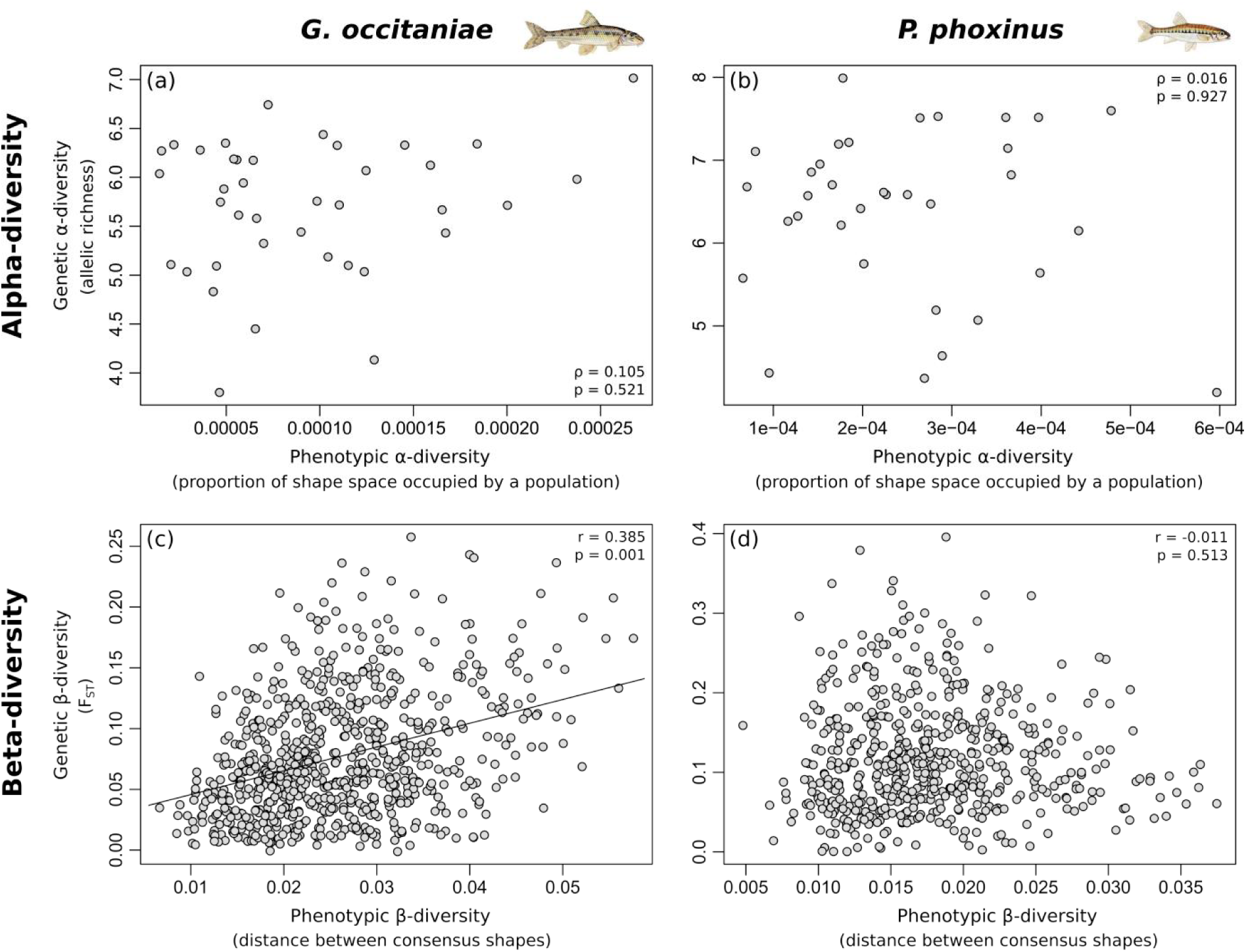
Genetic α-diversity (allelic richness) of Gobio occitaniae (a) and Phoxinus phoxinus (b) plotted against phenotypic α-diversity (proportion of shape space occupied by each population) with Spearman’s rho and associated P-values; and genetic β-diversity (FST) of Gobio occitaniae (c) and Phoxinus phoxinus (d) plotted against β-diversity (euclidean distance between the consensus shapes of each pair of populations) with Mantel’s r and associated P-values.

Mean between-sites genetic β-diversity was in average lower in *G. occitaniae* than in *P. phoxinus* (Wilcoxon rank sum test, W = 245, P < 0.001, Fig. 3c), whereas the reverse held true for mean phenotypic β-diversity per site (Wilcoxon rank sum test, W = 1284, P < 0.001, Fig. 3d); gudgeon populations were -in average-less genetically differentiated than minnow populations but more phenotypically differentiated. The fact that *G. occitaniae* populations were more phenotypically differentiated was confirmed using Fβ (Wilcoxon rank sum test, W = 1259, P < 0.001; Fig. S5). We further found that phenotypic turnover (measured as Fpturn) was also higher for *G. occitaniae* populations than for *P. phoxinus* populations (Wilcoxon rank sum test, W = 981, P < 0.001; Fig. S5). Remarkably, for 116 out of 741 populations pairs of gudgeon, Fβ and Fpturn were equal to 1, indicating no overlap in the portions of the shape space occupied by populations, whereas in *P. phoxinus*, none of the populations pair had values of Fβ and Fpturn equal to 1. The correlation between genetic and phenotypic β-diversity was positive and significant in *G. occitaniae* (i.e. significant β-GPIDC, Mantel test, r = 0.358, P = 0.001, Fig. 4c) but not in *P. phoxinus* (Mantel test, r = -0.011, P = 0.521, Fig. 4d).

### Determinants of α- and β-GPIDCs

*α-GPIDCs.* In *G. occitaniae* and *P. phoxinus*, the models with the lowest AIC scores were well supported by the data, as indicated by non-significant p-values associated with the tests of conditional independences (C = 75.485, d.f. = 74, P = 0.389 for *G. occitaniae* and C = 76.524, d.f. = 72, P = 0.335 for *P. phoxinus*, Table 2a). In *G. occitaniae*, we found a negative effect of *isolation* on both genetic and phenotypic α*-*diversity, indicating that populations were genetically and phenotypically impoverished in sites situated at high altitude and far from the river mouth. Additionally, phenotypic α*-*diversity tended to be negatively related to connectivity (Fig. 5a). In *P. phoxinus*, genetic α*-*diversity was also negatively related to *isolation*, but not phenotypic α*-*diversity (Fig. 5b). Genetic α*-*diversity was also positively related to habitat area. Phenotypic α*-*diversity was negatively correlated to oxygen saturation, which was in turn positively associated with *isolation* and habitat area (Fig. 5b).

**FIGURE 5.**
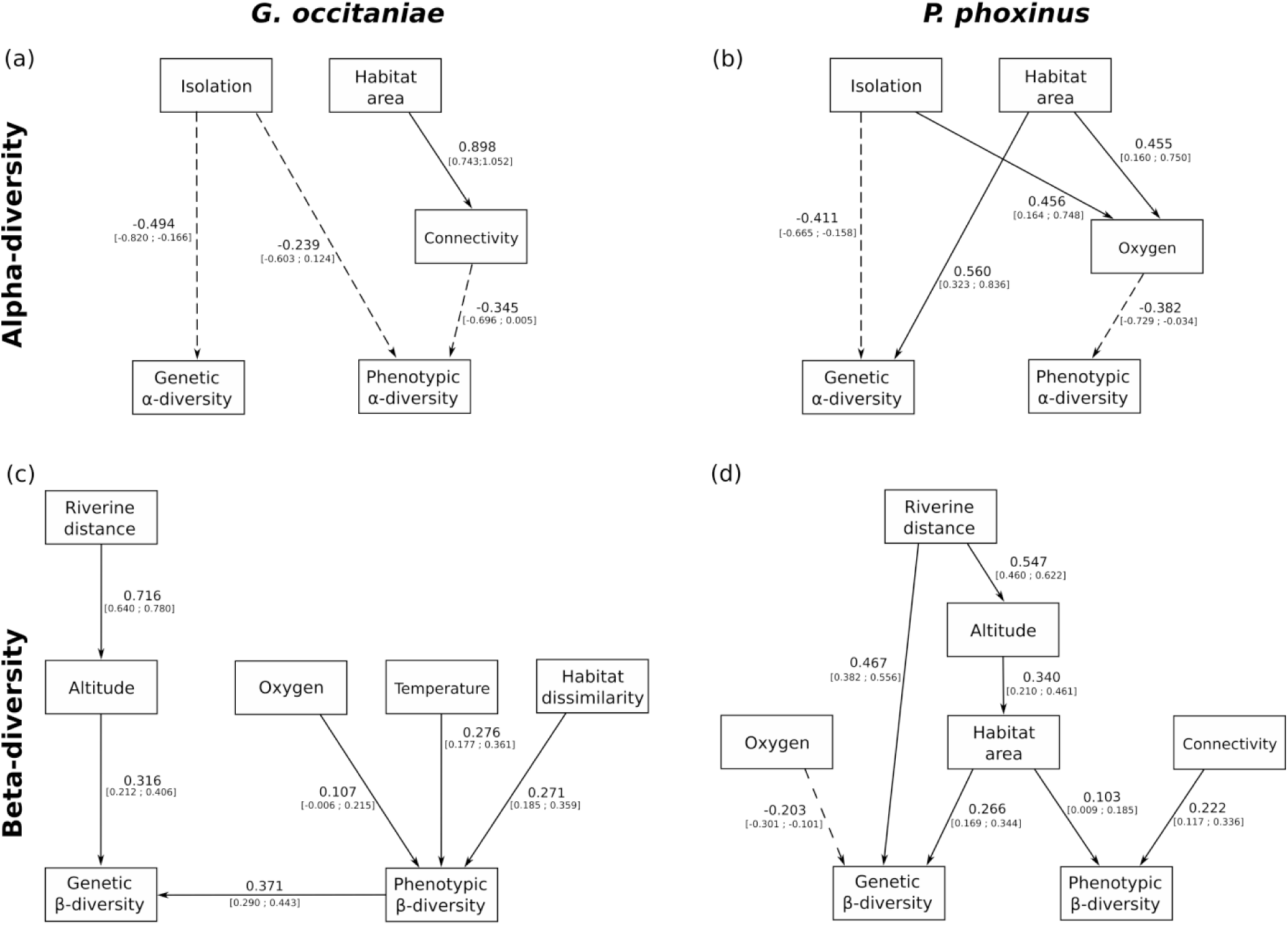
Graphical representations of the models describing the causal relationships between environmental variables and genetic and phenotypic α-diversity in Gobio occitaniae (a) and Phoxinus phoxinus (b), and between environmental variables and genetic and phenotypic β- diversity in Gobio occitaniae (a) and Phoxinus phoxinus (b), obtained using the d-sep test. Single-headed arrows indicate a causal path. Solid and dashed lines stand for positive and negative values, respectively.

**Table 2:** D-sep test statistics used to disentangle the effects of environmental variables on genetic and phenotypic α-diversity (a) and on genetic and phenotypic β-diversity (b) in *Gobio occitaniae* and *Phoxinus phoxinus*. For each species and diversity facet, we simplified a full model (i.e. a model including all paths described in the main text) until reaching the models with the lowest AIC score represented in Figure 5.

| **(a) Alpha-intraspecific diversity** | C statistics | d.f. | p-value | AIC |
| --- | --- | --- | --- | --- |
| *Gobio occitaniae* |  |  |  |  |
| Complete model | 32.569 | 30 | 0.342 | 128.569 |
| Optimal model | 75.485 | 74 | 0.430 | 116.791 |
| *Phoxinus phoxinus* |  |  |  |  |
| Complete model | 51.971 | 30 | 0.008 | 147.971 |
| Optimal model | 76.524 | 72 | 0.335 | 122.524 |
| **(b) Bêta-intraspecific diversity** | C statistics | d.f. | p-value | AIC |
| *Gobio occitaniae* |  |  |  |  |
| Complete model | 47.866 | 30 | 0.020 | 167.866 |
| Optimal model | 108.119 | 92 | 0.120 | 150.119 |
| *Phoxinus phoxinus* |  |  |  |  |
| Complete model | 50.105 | 30 | 0.012 | 170.105 |
| Optimal model | 97.058 | 94 | 0.394 | 133.058 |

### β-GPIDCs

The models with the lowest AIC scores were well supported by the data in both species (C = 109.167, 92 d.f., P = 0.130 in *G. occitaniae* and C = 97.699, 94 d.f., P = 0.575 in *P. phoxinus*, Table 2b). For *G. occitaniae*, genetic β-diversity was positively related to the cumulative difference in altitude, which was itself related to riverine distance (leading to an indirect relationship between genetic β-diversity and riverine distance, Fig. 5c). Phenotypic β-diversity was positively correlated to three environmental variables (difference in oxygen concentration, difference in water temperature and habitat dissimilarity, Fig. 5d). Additionally, we found a positive relationship between genetic and phenotypic β-diversity (Fig. 5d). Regarding *P. phoxinus*, genetic β-diversity was positively related to riverine distance both directly and indirectly through difference in altitude and difference in habitat area. Genetic β-diversity was also negatively related to difference in oxygen. Phenotypic β- diversity was directly related to difference in connectivity and indirectly related to pairwise riverine distance through difference in altitude and habitat area (Fig. 5d).

### Phenotype-environment relationships

In *G. occitaniae*, the first relative warp had high values in individuals living in sites with high concentration in oxygen (β = 0.272, CI = [0.138; 0.406]) and low water temperature (β = -0.236, CI = [-0.367; -0.105]), and where the proportion of silt in the substrate was low (β = -0.268, CI = [-0.401; -0.136]) (Table 3). Additionally, the first relative warp was related to individual centroid size (used as a proxy for individual size) and its quadratic term (β = 0.513, CI = [0.469; 0.557] and β = -0.106, CI = [-0.136; -0.076] respectively), suggesting allometric relationships with this relative warp. None of the environmental variables we considered were likely to be associated to the second relative warp. In *P. phoxinus*, the first and second relative warps were significantly associated to the individual centroid size (β = 0.455, CI = [0.392; 0.518] and β = -0.342, CI = [-0.413; -0.271] respectively), which also suggests allometric relationships. Surprisingly, we found no phenotype-environment relationships in *P. phoxinus*, suggesting that, in this species, most of the phenotype variations we detected were independent of environmental characteristics and topography.

## Discussion

In this study, we tested for Genetic-Phenotypic Intraspecific Diversity correlations (GPIDCs) in two parapatric freshwater fish species, and explored the processes shaping the spatial distribution of their genetic and phenotypic characteristics. Our results revealed disparities in the distribution of genetic and phenotypic diversity in the two species, as well as common and contrasted processes shaping diversity at the α- and β-levels.

In terms of genetic diversity, we found that, overall, *P. phoxinus* populations were more locally diverse (higher neutral genetic α-diversity) and more differentiated (higher neutral genetic β-diversity) than *G. occitaniae* populations (Fig. 3a and 3c). The overall higher genetic α-diversity in *P. phoxinus* may indicate different evolutionary histories between the two species. For instance, this may indicate that ancient effective population sizes were higher and more stable over time in *P. phoxinus* than in *G. occitaniae*, thus limiting the impact of drift, and/or that multiple glacial refugia existed in *P. phoxinus*, hence favouring the maintenance of a high neutral genetic diversity (Hewitt 1999). The overall high genetic α- diversity found in *P. phoxinus* populations may also result from high levels of biological connectivity in this species when compared to *G. occitaniae* (Frankham 1996) as suggested by spatial patterns of isolation-by-distance (Appendix SX). We also found that local levels of neutral genetic α-diversity in *P. phoxinus* were lower in sites of small habitat area (Fig. 5a), indicating an increased effect of genetic drift in habitat with small carrying capacity. This relationship between neutral genetic α-diversity and habitat area was not observed in *G. occitaniae*, probably because of a sampling bias, as *G. occitaniae* was generally found at lower altitudes and thus in stretches of higher carrying capacity than *P. phoxinus* (mean habitat area: 29.9 m and 16.2 m in *G. occitaniae* and *P. phoxinus* respectively; anova on log-transformed data: F = 4.16, df = 71, p = 0.06). Higher overall levels of neutral genetic β-diversity in *P. phoxinus* (Fig. 3c) may also be explained by this possible altitudinal sampling bias, responsible for higher spatial heterogeneity in the influence of drift (Prunier et al. 2017). Accordingly, we observed a positive impact of differences in habitat areas on genetic β-diversity in *P. phoxinus* (Fig. 5d), indicating that -as expected-populations experiencing contrasted intensity of genetic drift were more genetically differentiated (Prunier et al. 2017).

Despite these differences, we found common processes driving genetic diversity in both species. First, we found that neutral genetic α-diversity was strongly related to geographic isolation in both species, with lower genetic diversity observed in highly isolated sites, i.e. sites at high altitude and far from the river mouth (Fig. 5a-b). This decrease in neutral genetic diversity in geographically isolated sites has already been reported, and has actually been suggested to be a general pattern in riverine networks (Paz-Vinas et al. 2015). Two non-exclusive hypotheses can explain this pattern. First, movements between populations might be directionally-biased due to water flow (Morrissey and de Kerckhove 2009, Paz-Vinas et al. 2013). This asymmetric dispersal leads to an increase in gene flow from upstream (isolated sites) to downstream, generating an upstream loss of genetic diversity through emigration (Kawecki and Holt 2002). Alternatively, a decrease in genetic diversity in upstream sites might reflect the species colonization history from downstream glacial refugia. Second, genetic β-diversity was driven by topographic features in both species (Fig. 5a-b); in 1. *G. occitaniae*, genetic differentiation was higher between sites isolated from each other by high altitude drops along the network, whereas in *P. phoxinus*, genetic differentiation was higher between sites separated by a high riverine distance. These two latter patterns confirm the existence of a process of isolation-by-distance (Hutchison and Templeton 1999) in the two species (supplementary material Appendix SX).

At the phenotypic level, our findings suggest that the regional pool in *G. occitaniae* was composed of poorly diverse local populations (low phenotypic α-diversity; Fig. 3b) that were highly dissimilar from one site to another (high phenotypic β-diversity with high turnover between populations, i.e. different populations display different phenotypes; Fig. 3d). Conversely, in *P. phoxinus*, phenotypic α-diversity was higher and phenotypic β-diversity was lower than in *G. occitaniae*, which suggests that the regional pool of *P. phoxinus* was composed of highly diverse local populations that were highly similar from one site to another. The contrasted morphological patterns found in these two parapatric species may result (i) from higher effective population sizes in *P. phoxinus* than in *G. occitaniae* (as suggested by measures of genetic α-diversity, see above), and/or (ii) from stronger effects of selection (or environmental effects in general) in *G. occitaniae* than in *P. phoxinus*. Indeed, a stronger effect of selection is expected to lead to environmental filtering and hence to less phenotypically diverse populations at the local scale (local adaptation) as well as to a high phenotypic β-diversity between populations resulting from adaptive divergence (and/or strong plastic effects) (Blanquart et al. 2013). This later hypothesis was strengthened by the significant relationships found between the individual shapes in *G. occitaniae* and three environmental variables (see below), and by the limited scale of gene flow in *G. occitaniae* when compared to isolation-by-distance pattern in *P. phoxinus* (supplementary material Appendix SX). Despite the strong environmental heterogeneity measured among sites, *P. phoxinus* populations appeared highly similar suggesting a higher level of generalism in *P. phoxinus* populations than in possibly more specialist *G. occitaniae* populations, as well as a homogenizing influence of effective dispersal in *P. phoxinus*.

In line with this result, we found highly contrasted processes shaping phenotypic diversity in both species. In *G. occitaniae*, phenotypic α-diversity was lower in highly-connected sites, with a high centrality index (Fig. 5a). This result was unexpected since highly central sites are expected to receive more dispersers, hence enhancing phenotypic diversity and impeding local adaptation. However, the observed pattern could be explained by a higher efficiency of selection in central sites in which dispersal introduces additional phenotype variability necessary for adaptation (Lenormand 2002), potentially in combination with a habitat matching process, that would hinder the negative impact of gene flow on local adaptation (Edelaar et al. 2008). Alternatively -and not-exclusively-this negative relationship could arise from a statistical bias, for instance if an unmeasured collinear variable explained both centrality and phenotypic α-diversity. However, phenotypic α-diversity tended to be lower in isolated sites (in which, according to our former hypothesis, populations are expected to be less locally adapted and hence more diverse), which may suggest an effect of neutral processes (“phenotypic” drift) as observed in neutral genetic diversity, and/or stronger effects of environmental filtering in isolated sites (high altitude and far from the river mouth) than in less isolated sites. This latter hypothesis of strong environmental filtering is likely given that upstream (isolated) sites are known to experience harsh environmental conditions (Vannote et al. 1980). Furthermore, in *G. occitaniae*, phenotypic β-diversity was primarily shaped by environmental variables related to habitat and water features (namely, difference in oxygen saturation, temperature and habitat dissimilarity) such that mean phenotype was different between sites displaying contrasted abiotic conditions (Fig. 5c). This impact of environment on phenotype was strengthened by the direct relationships found between individual phenotype and oxygen saturation, temperature and proportion of silt in the habitat. These two results confirm the hypothesis that selection (or environment in general) has strong effects on phenotype in *G. occitaniae*, however it remains unclear whether these effects originate from heritable differentiation or environmentally induced plasticity.

In *P. phoxinus*, phenotypic α-diversity was higher in sites with low oxygen concentration, suggesting a positive influence of stressful conditions on phenotypic α-diversity (Fig. 5b). This result was surprising as we expected that a low saturation in oxygen would sustain small population sizes, hence reducing phenotypic diversity. Moreover, stressful conditions were expected to strengthen selection pressure. However, stressful conditions have already been proven to have a positive effect on intraspecific diversity, notably (i) when they lead to an increase of mutation and recombination rates in non-neutral parts of the genome (Badyaev 2005), and (ii) when they lead to an increase in phenotypic plasticity (Ghalambor et al. 2007, Rey et al. 2016). Phenotypic β-diversity was increased between populations inhabiting sites of different area and different connectivity (Fig. 5d). These relations may suggest an effect of neutral processes associated with population sizes and gene flow on phenotypic diversity, which is likely as local adaptation does not appear to be high in this species.

We found no GPIDCs at the α-level in either species, indicating that neutral genetic α-diversity and phenotypic α-diversity are driven by independent processes. Although consistent with our theoretical expectations, this result was surprising in *G. occitaniae* as we found a similar impact of isolation on genetic and phenotypic α-diversity. This absence of correlation suggests that the influence of other processes (related to connectivity) were strong enough to break spatial covariation between these two facets of diversity in this species.

At the β-level, we found a significant and positive GPIDC in *G. occitaniae*, such that populations being genetically different were also phenotypically different. However, this correlation did not seem to originate from similar environmental processes shaping both facets of β-diversity but appeared to be mainly caused by a direct effect of one facet of β-diversity on the other. It was not possible to statistically determine the direction of this relation (genetic diversity to phenotypic diversity or phenotypic diversity to genetic diversity) due to methodological limitations. However, given that we focused on neutral genetic markers, a direct impact of genetic β-diversity on phenotypic β-diversity seems unlikely except if we assume (i) that the here chosen microsatellite markers properly reflect the genomic diversity in this species and (ii) that phenotypic diversity in this species is mainly driven by the genetic background of individuals. Alternatively, positive assortative mating (i.e. the propensity to mate with phenotypically similar individuals) has been shown to be particularly strong in fish (Jiang et al. 2013) and could explain this direct relation between phenotypic and genetic differentiation (Wang and Summers 2010). Yet, although our dataset encompasses the main environmental variables known to be involved in adaptive and neutral processes in freshwater fish, we cannot rule out the influence of a possible unmeasured abiotic or biotic factor impacting both facets of β-diversity. In *P. phoxinus*, genetic and phenotypic β-diversity were not correlated despite of a similar impact of habitat area on both facets of diversity. Other important processes involving riverine distance and connectivity could impede spatial covariation between these two facets of diversity in this species.

The use of an integrative framework allowed us to unveil striking dissimilarities between the patterns and drivers of genetic and phenotypic intraspecific diversity in two parapatric freshwater fish species. First, we found indications of limited gene flow and of local adaptation in *G. occitaniae* populations. Second, we observed that, in *P. phoxinus*, populations were phenotypically more diverse and that gene flow occurred at a larger spatial scale. This high phenotypic diversity could indicate a bet-hedging strategy (i.e. the augmentation of phenotypic diversity to optimize fitness in varying environments), possibly in response to inter-annual variation in local flow regimes (Lytle and Poff 2004). Studying neutral genetic diversity and phenotypic diversity within an integrative framework hence appeared as a valuable way of deciphering the complex and diverse impacts of neutral and adaptive processes on intraspecific diversity.

While introducing the novel framework of Species-Genetic Diversity Correlation, Vellend (2005) stated that treating interspecific and intraspecific diversity as independent phenomena in community ecology and population genetics was irrelevant. Similarly, genetic and phenotypic diversity are clearly interrelated but are mainly studied separately in population genetics and functional ecology. We advocate for a greater integration across disciplinary boundaries in future studies in order to advance our understanding of the distribution of intraspecific diversity.

## References

Adams, D. C., and E. Otárola-Castillo. 2013. geomorph: an R package for the collection and analysis of geometric morphometric shape data. Methods in Ecology and Evolution 4:393–399.

Aljanabi, S. M., and I. Martinez. 1997. Universal and rapid salt-extraction of high quality genomic DNA for PCR-based techniques. Nucleic acids research 25:4692–4693.

Badyaev, A. V. 2005. Stress-induced variation in evolution: from behavioural plasticity to genetic assimilation. Proceedings of the Royal Society of London B: Biological Sciences 272:877–886.

Bartoń, K. 2016. MuMIn: Multi-Model Inference.

Baselga, A. 2010. Partitioning the turnover and nestedness components of beta diversity. Global Ecology and Biogeography 19:134–143.

Blanchet, S., J. G. Prunier, and H. De Kort. 2017. Time to Go Bigger: Emerging Patterns in Macrogenetics. Trends in Genetics 33:579–580.

Blanquart, F., O. Kaltz, S. L. Nuismer, and S. Gandon. 2013. A practical guide to measuring local adaptation. Ecology Letters 16:1195–1205.

Bolnick, D. I., P. Amarasekare, M. S. Araújo, R. Bürger, J. M. Levine, M. Novak, V. H. W. Rudolf, S. J. Schreiber, M. C. Urban, and D. A. Vasseur. 2011. Why intraspecific trait variation matters in community ecology. Trends in Ecology & Evolution 26:183–192.

Bolnick, D. I., R. Svanbäck, J. A. Fordyce, L. H. Yang, J. M. Davis, C. D. Hulsey, and M. L. Forister. 2003. The ecology of individuals: incidence and implications of individual specialization. The American Naturalist 161:1–28.

Bookstein, F. L. 1991. Morphometric Tools for Landmark Data: Geometry and Biology. Cambridge University Press, Cambridge ; New York, NY.

Buisson, L., L. Blanc, and G. Grenouillet. 2008. Modelling stream fish species distribution in a river network: the relative effects of temperature versus physical factors. Ecology of Freshwater Fish 17:244–257.

Burnham, K. P., and D. R. Anderson. 2002. Model Selection and Multimodel Inference: A Practical Information-Theoretic Approach. 2nd edition. Springer Verlag, New York, NY.

Caschili, S. 2010. ComplexNetGIS: a tool for the analysis of complex spatial networks. Pages 233–242 *in* G. Las Casas, P. Pontrandolfi, and B. Murgante, editors. Informatica e pianificazione urbana e territoriale. Libria, Melfi.

Chave, J. 2013. The problem of pattern and scale in ecology: what have we learned in 20 years? Ecology Letters 16:4–16.

Crispo, E., and L. J. Chapman. 2008. Population genetic structure across dissolved oxygen regimes in an African cichlid fish. Molecular Ecology 17:2134–2148.

Dray, S., and A.-B. Dufour. 2007. The ade4 Package: Implementing the Duality Diagram for Ecologists. Journal of Statistical Software 22:1–20.

Edelaar, P., A. M. Siepielski, and J. Clobert. 2008. Matching Habitat Choice Causes Directed Gene Flow: A Neglected Dimension in Evolution and Ecology. Evolution 62:2462– 2472.

Estrada, E., and Ö. Bodin. 2008. Using network centrality measures to manage landscape connectivity. Ecological Applications 18:1810–1825.

Fourtune, L., I. Paz-Vinas, G. Loot, J. G. Prunier, and S. Blanchet. 2016. Lessons from the fish: a multi-species analysis reveals common processes underlying similar species-genetic diversity correlations. Freshwater Biology 61:1830–1845.

Fourtune, L., J. G. Prunier, I. Paz-Vinas, G. Loot, C. Veyssière, and S. Blanchet. 2018. Inferring Causalities in Landscape Genetics: An Extension of Wright’s Causal Modeling to Distance Matrices. The American Naturalist:491–508.

Frankham, R. 1996. Relationship of Genetic Variation to Population Size in Wildlife. Conservation Biology 10:1500–1508.

Freeman, L. 1977. A Set of Measures of Centrality Based on Betweenness. Sociometry 40:35–41.

Fronhofer, E. A., and F. Altermatt. 2017. Classical metapopulation dynamics and eco-evolutionary feedbacks in dendritic networks. Ecography.

Ghalambor, C. K., J. K. McKAY, S. P. Carroll, and D. N. Reznick. 2007. Adaptive versus non-adaptive phenotypic plasticity and the potential for contemporary adaptation in new environments. Functional Ecology 21:394–407.

Grant, E. H. C., W. H. Lowe, and W. F. Fagan. 2007. Living in the branches: population dynamics and ecological processes in dendritic networks. Ecology Letters 10:165– 175.

Hartl, D. L., and A. G. Clark. 2007. Principles of population genetics. 4th edition. Sinauer Associates, Sunderland, MA.

Hedrick, P. W. 2006. Genetic Polymorphism in Heterogeneous Environments: The Age of Genomics. Annual Review of Ecology, Evolution, and Systematics 37:67–93.

Hewitt, G. M. 1999. Post-glacial re-colonization of European biota. Biological Journal of the Linnean Society 68:87–112.

Hoffman, J. I., F. Simpson, P. David, J. M. Rijks, T. Kuiken, M. A. S. Thorne, R. C. Lacy, and K. K. Dasmahapatra. 2014. High-throughput sequencing reveals inbreeding depression in a natural population. Proceedings of the National Academy of Sciences 111:3775–3780.

Holderegger, R., U. Kamm, and F. Gugerli. 2006. Adaptive vs. neutral genetic diversity: implications for landscape genetics. Landscape Ecology 21:797–807.

Hughes, A. R., B. D. Inouye, M. T. J. Johnson, N. Underwood, and M. Vellend. 2008. Ecological consequences of genetic diversity. Ecology Letters 11:609–623.

Hutchison, D., and A. Templeton. 1999. Correlation of pairwise genetic and geographic distance measure: inferring the relative influences of gene flow and drift on distribution of genetic variability. Evolution 53:1898–1914.

Jiang, Y., D. I. Bolnick, and M. Kirkpatrick. 2013. Assortative Mating in Animals. The American Naturalist 181:E125–E138.

Jost, L. 2008. GST and its relatives do not measure differentiation. Molecular Ecology 17:4015–4026.

Jung, V., C. H. Albert, C. Violle, G. Kunstler, G. Loucougaray, and T. Spiegelberger. 2013. Intraspecific trait variability mediates the response of subalpine grassland communities to extreme drought events. Journal of Ecology 102:45–53.

Kawecki, T. J., and D. Ebert. 2004. Conceptual issues in local adaptation. Ecology Letters 7:1225–1241.

Kawecki, T. J., and R. D. Holt. 2002. Evolutionary consequences of asymmetric dispersal rates. The American Naturalist 160:333–347.

Lamy, T., F. Laroche, P. David, F. Massol, and P. Jarne. 2017. The contribution of species– genetic diversity correlations to the understanding of community assembly rules. Oikos 126:759–771.

Leimar, O. 2005. The Evolution of Phenotypic Polymorphism: Randomized Strategies versus Evolutionary Branching. The American Naturalist 165:669–681.

Leinonen, T., R. J. S. McCairns, R. B. O’Hara, and J. Merilä. 2013. QST–FST comparisons: evolutionary and ecological insights from genomic heterogeneity. Nature Reviews Genetics 14:179–190.

Lenormand, T. 2002. Gene flow and the limits to natural selection. Trends in Ecology & Evolution 17:183–189.

Loreau, M. 2000. Biodiversity and ecosystem functioning: recent theoretical advances. Oikos 91:3–17.

Lowe, W. H., R. P. Kovach, and F. W. Allendorf. 2017. Population Genetics and Demography Unite Ecology and Evolution. Trends in Ecology & Evolution 32:141– 152.

Lowe, W. H., and M. A. McPeek. 2014. Is dispersal neutral? Trends in Ecology & Evolution 29:444–450.

Lytle, D. A., and N. L. Poff. 2004. Adaptation to natural flow regimes. Trends in Ecology & Evolution 19:94–100.

Manel, S., M. K. Schwartz, G. Luikart, and P. Taberlet. 2003. Landscape genetics: combining landscape ecology and population genetics. Trends in Ecology & Evolution 18:189– 197.

Mimura, M., T. Yahara, D. P. Faith, E. Vázquez-Domínguez, R. I. Colautti, H. Araki, F. Javadi, J. Núñez-Farfán, A. S. Mori, S. Zhou, P. M. Hollingsworth, L. E. Neaves, Y. Fukano, G. F. Smith, Y.-I. Sato, H. Tachida, and A. P. Hendry. 2017. Understanding and monitoring the consequences of human impacts on intraspecific variation. Evolutionary Applications 10:121–139.

Moran, E. V., F. Hartig, and D. M. Bell. 2015. Intraspecific trait variation across scales: implications for understanding global change responses. Global Change Biology 22:137–150.

Morrissey, M. B., and D. T. de Kerckhove. 2009. The Maintenance of Genetic Variation Due to Asymmetric Gene Flow in Dendritic Metapopulations. The American Naturalist 174:875–889.

Nei, M., F. Tajima, and Y. Tateno. 1983. Accuracy of estimated phylogenetic trees from molecular data. Journal of Molecular Evolution 19:153–170.

Odling-Smee, F. J., K. N. Laland, and M. W. Feldman. 2003. Niche Construction: The Neglected Process in Evolution. Princeton University Press, Princeton, NJ and Oxford.

Outomuro, D., and F. Johansson. 2017. A potential pitfall in studies of biological shape: does size matter? Journal of Animal Ecology 86:1447–1457.

Paz-Vinas, I., and S. Blanchet. 2015. Dendritic connectivity shapes spatial patterns of genetic diversity: a simulation-based study. Journal of Evolutionary Biology 28:986–994.

Paz-Vinas, I., G. Loot, V. M. Stevens, and S. Blanchet. 2015. Evolutionary processes driving spatial patterns of intra-specific genetic diversity in river ecosystems. Molecular Ecology 24:4586–4604.

Paz-Vinas, I., E. Quéméré, L. Chikhi, G. Loot, and S. Blanchet. 2013. The demographic history of populations experiencing asymmetric gene flow: combining simulated and empirical data. Molecular Ecology 22:3279–3291.

Pearl, J., and T. Verma. 1987. The logic of representing dependencies by directed graphs. Pages 374–379 Proceedings of the sixth National conference on Artificial intelligence - Volume 1. AAAI Press, Seattle, WA.

Pella, H., J. Lejot, N. Lamouroux, and T. Snelder. 2012. Le réseau hydrographique théorique (RHT) français et ses attributs environnementaux. Géomorphologie : relief, processus, environnement 3:317–336.

Pinheiro, J., D. Bates, S. Debroy, and D. Sarkar. 2016. nlme: Linear and Nonlinear Mixed Effects Models.

Prunier, J. G., V. Dubut, L. Chikhi, and S. Blanchet. 2017. Contribution of spatial heterogeneity in effective population sizes to the variance in pairwise measures of genetic differentiation. Methods in Ecology and Evolution Early view.

Quantum GIS Development Team. 2017. Quantum GIS Geographic Information System. Open Source Geospatial Foundation Project.

R Development Core Team. 2017. R: A language and environment for statistical computing. R Foundation for Statistical Computing, Vienna, Austria.

Raeymaekers, J. A. M., G. E. Maes, S. Geldof, I. Hontis, K. Nackaerts, and F. A. M. Volckaert. 2008. Modeling genetic connectivity in sticklebacks as a guideline for river restoration. Evolutionary Applications 1:475–488.

Reed, D. H., and R. Frankham. 2003. Correlation between Fitness and Genetic Diversity. Conservation Biology 17:230–237.

Relethford, J. H. 1991. Genetic Drift and Anthropometric Variation in Ireland. Human Biology 63:155–165.

Rey, O., E. Danchin, M. Mirouze, C. Loot, and S. Blanchet. 2016. Adaptation to Global Change: A Transposable Element–Epigenetics Perspective. Trends in Ecology & Evolution 31:514–526.

Rohlf, F. J. 1993. Relative warp analysis and an example of its application to mosquito wings. Pages 131–160 *in* L. F. Marcus, E. Bello, and A. Garcia-Valdecasas, editors. Contribution to Morphometrics. Museo Nacional de Ciencias Naturales, Consejo Superior de Investigaciones Científicas, CSIC, Madrid.

Rohlf, F. J., and L. F. Marcus. 1993. A revolution morphometrics. Trends in Ecology & Evolution 8:129–132.

Rohlf, F. J., and D. Slice. 1990. Extensions of the Procrustes Method for the Optimal Superimposition of Landmarks. Systematic Biology 39:40–59.

Rousset, F. 1997. Genetic Differentiation and Estimation of Gene Flow from F-Statistics Under Isolation by Distance. Genetics 145:1219.

Rousset, F. 2008. genepop’007: a complete re-implementation of the genepop software for Windows and Linux. Molecular Ecology Resources 8:103–106.

Schneider, C. A., W. S. Rasband, and K. W. Eliceiri. 2012. NIH Image to ImageJ: 25 years of image analysis. Nature Methods 9:671–675.

Sexton, J. P., S. B. Hangartner, and A. A. Hoffmann. 2014. Genetic Isolation by Environment or Distance: Which Pattern of Gene Flow Is Most Common? Evolution 68:1–15.

Shipley, B. 2000. A new inferential test for path models based on directed acyclic graphs. Structural Equation Modeling 7:206–218.

Shipley, B. 2013. The AIC model selection method applied to path analytic models compared using a d-separation test. Ecology 94:560–564.

Siefert, A., C. Violle, L. Chalmandrier, C. H. Albert, A. Taudiere, A. Fajardo, L. W. Aarssen, C. Baraloto, M. B. Carlucci, M. V. Cianciaruso, V. de L. Dantas, F. de Bello, L. D. S. Duarte, C. R. Fonseca, G. T. Freschet, S. Gaucherand, N. Gross, K. Hikosaka, B. Jackson, V. Jung, C. Kamiyama, M. Katabuchi, S. W. Kembel, E. Kichenin, N. J. B. Kraft, A. Lagerström, Y. L. Bagousse-Pinguet, Y. Li, N. Mason, J. Messier, T. Nakashizuka, J. M. Overton, D. A. Peltzer, I. M. Pérez-Ramos, V. D. Pillar, H. C. Prentice, S. Richardson, T. Sasaki, B. S. Schamp, C. Schöb, B. Shipley, M. Sundqvist, M. T. Sykes, M. Vandewalle, and D. A. Wardle. 2015. A global meta-analysis of the relative extent of intraspecific trait variation in plant communities. Ecology Letters 18:1406–1419.

Szpiech, Z. A., M. Jakobsson, and N. A. Rosenberg. 2008. ADZE: a rarefaction approach for counting alleles private to combinations of populations. Bioinformatics 24:2498–2504.

Taberlet, P., N. E. Zimmermann, T. Englisch, A. Tribsch, R. Holderegger, N. Alvarez, H. Niklfeld, G. Coldea, Z. Mirek, A. Moilanen, W. Ahlmer, P. A. Marsan, E. Bona, M. Bovio, P. Choler, E. Cieślak, L. Colli, V. Cristea, J.-P. Dalmas, B. Frajman, L. Garraud, M. Gaudeul, L. Gielly, W. Gutermann, N. Jogan, A. A. Kagalo, G. Korbecka, P. Küpfer, B. Lequette, D. R. Letz, S. Manel, G. Mansion, K. Marhold, F. Martini, R. Negrini, F. Niño, O. Paun, M. Pellecchia, G. Perico, H. Piękoś-Mirkowa, F. Prosser, M. Puşcaş, M. Ronikier, M. Scheuerer, G. M. Schneeweiss, P. Schönswetter, L. Schratt-Ehrendorfer, F. Schüpfer, A. Selvaggi, K. Steinmann, C. Thiel-Egenter, M. van Loo, M. Winkler, T. Wohlgemuth, T. Wraber, F. Gugerli, and IntraBioDiv Consortium. 2012. Genetic diversity in widespread species is not congruent with species richness in alpine plant communities. Ecology Letters 15:1439–1448.

Van Oosterhout, C., W. F. Hutchinson, D. P. M. Wills, and P. Shipley. 2004. micro-checker: software for identifying and correcting genotyping errors in microsatellite data. Molecular Ecology Notes 4:535–538.

Vannote, R. L., G. W. Minshall, K. W. Cummins, J. R. Sedell, and C. E. Cushing. 1980. The River Continuum Concept. Canadian Journal of Fisheries and Aquatic Sciences 37:130–137.

Vellend, M. 2005. Species diversity and genetic diversity: parallel processes and correlated patterns. The American Naturalist 166:199–215.

Vellend, M., G. Lajoie, A. Bourret, C. Múrria, S. W. Kembel, and D. Garant. 2014. Drawing ecological inferences from coincident patterns of population- and community-level biodiversity. Molecular Ecology 23:2890–2901.

Villéger, S., G. Grenouillet, and S. Brosse. 2013. Decomposing functional β-diversity reveals that low functional β-diversity is driven by low functional turnover in European fish assemblages: Decomposing functional β-diversity. Global Ecology and Biogeography 22:671–681.

Villéger, S., N. W. H. Mason, and D. Mouillot. 2008. New multidimensional functional diversity indices for a multifaceted framework in functional ecology. Ecology 89:2290–2301.

Villéger, S., P. M. Novack-Gottshall, and D. Mouillot. 2011. The multidimensionality of the niche reveals functional diversity changes in benthic marine biotas across geological time. Ecology Letters 14:561–568.

Violle, C., B. J. Enquist, B. J. McGill, L. Jiang, C. H. Albert, C. Hulshof, V. Jung, and J. Messier. 2012. The return of the variance: intraspecific variability in community ecology. Trends in Ecology & Evolution 27:244–252.

Violle, C., M.-L. Navas, D. Vile, E. Kazakou, C. Fortunel, I. Hummel, and E. Garnier. 2007. Let the concept of trait be functional! Oikos 116:882–892.

Wang, I. J., and G. S. Bradburd. 2014. Isolation by environment. Molecular Ecology 23:5649–5662.

Wang, I. J., and K. Summers. 2010. Genetic structure is correlated with phenotypic divergence rather than geographic isolation in the highly polymorphic strawberry poison-dart frog. Molecular Ecology 19:447–458.

